# An unusual amino acid substitution within hummingbird cytochrome *c* oxidase alters a key proton-conducting channel

**DOI:** 10.1101/610915

**Authors:** Cory D. Dunn, Bala Anı Akpınar, Vivek Sharma

## Abstract

Hummingbirds in flight exhibit the highest metabolic rate of all vertebrates. The bioenergetic requirements associated with sustained hovering flight raise the possibility of unique amino acid substitutions that would enhance aerobic metabolism. Here, we have identified a non-conservative substitution within the mitochondria-encoded cytochrome *c* oxidase subunit I (COI) that is fixed within hummingbirds, but not among other vertebrates. This unusual change is also rare among metazoans, but can be identified in several clades with diverse life histories. We performed atomistic molecular dynamics simulations using bovine and hummingbird COI models, thereby bypassing experimental limitations imposed by the inability to modify mtDNA in a site-specific manner. Intriguingly, our findings suggest that COI amino acid position 153 (bovine numbering system) provides control over the hydration and activity of a key proton channel in COX. We discuss potential phenotypic outcomes linked to this alteration encoded by the hummingbird mitochondrial genome.

## INTRODUCTION

Hummingbirds are distinguished by their use of sustained hovering flight when feeding upon nectar and insects, when defending their territories, and when carrying out courtship displays (Hainsworth and Wolf 1972; Norberg 1996; Altshuler and Dudley 2002). Their exceptional mobility demands a prodigious level of mitochondrial ATP synthesis, and indeed, the metabolic rate of hummingbird flight muscles is exceedingly high (Lasiewski 1962; Suarez *et al.* 1991; Clark and Dudley 2010; Fernández *et al.* 2011). Many physiological and cellular features of hummingbirds appear to be tailored to their extreme metabolism, which can be maintained even within hypoxic environments up to 5000 meters above sea level (Projecto-Garcia *et al.* 2013). For example, hemoglobin structure (Projecto-Garcia *et al.* 2013) and cellular myoglobin concentration (Johansen *et al.* 1987) appear to be adapted to the oxygen delivery needs of hummingbirds. Additionally, the hearts of hummingbirds are larger, relative to their body size, than other birds and can pump at a rate of more than 1000 beats per minute (Bishop 1997). Beyond ATP synthesis, the metabolism of these tiny endotherms must also buffer against heat loss (Lasiewski 1963; López-Calleja and Bozinovic 1995; Suarez and Gass 2002). At the subcellular level, adaptation to the need for increased ATP and heat production can be readily visualized, since mitochondria in hummingbird flight muscles are highly, perhaps maximally, packed with cristae and are found in close apposition to capillaries (Suarez *et al.* 1991; Mathieu-Costello *et al.* 1992). Hummingbirds display an unexpectedly long lifespan when considering the allometric relationship between body mass and longevity (Calder 1990), but whether the hummingbird lifespan is linked to its unusual metabolic prowess is unclear.

Within the mitochondrial inner membrane, electrons progress through the electron transport chain (ETC), reach the cytochrome *c* oxidase (COX) complex, and are then used to reduce oxygen. Proton movements coupled to electron passage through COX contribute to the proton motive force (PMF) used for ATP production and thermogenesis (Wikström *et al.* 2018; Wikström and Sharma 2018). While several COX subunits are nucleus-encoded and imported to mitochondria, the core, catalytic subunits of COX (subunits COI, COII, and COIII) are encoded by mitochondrial DNA (mtDNA) (Johnston and Williams 2016), raising the possibility that unusual changes to the mitochondrial genome may have contributed to the remarkable metabolic properties of hummingbirds. Here, we identify an amino acid substitution in COI that is universal among hummingbirds, rare and unfixed among other birds and vertebrates, and limited to a small set of disparate clades among metazoans. Atomistic molecular dynamics (MD) simulations suggest that this substitution could affect COX function and may contribute to the uncommon physiological capabilities of hummingbirds.

## MATERIALS AND METHODS

### Sequence acquisition, alignment, phylogenetic analysis, and annotation

Mitochondrial proteomes were downloaded from the NCBI RefSeq database (O’Leary *et al.* 2016). Taxonomy analysis was performed using the ‘taxize’ package (Chamberlain and Szöcs 2013) and the NCBI taxonomy database (Federhen 2012), with manual curation when required. Beyond COI sequences acquired from the RefSeq database, additional COI barcodes were retrieved from the BOLD server (Ratnasingham and Hebert 2007).

Alignments were performed by use of standalone MAFFT (version 7.407) (Katoh and Standley 2013) or by T-coffee (version 13.40.5) in regressive mode (Garriga *et al.* 2019). For initial alignments of insect COI barcodes, MAFFT alignment was performed using an online server (Kuraku *et al.* 2013; Katoh 2017), and translations of barcodes using the appropriate codon tables were performed using AliView (Larsson 2014).

To seek mutations that occurred along the lineage to hummingbirds and to measure the relative conservation of the relative positions, a maximum-likelihood phylogenetic tree based upon an alignment of concatenated sequences of mtDNA-encoded proteins from the RefSeq database (release 92) was generated by FastTreeMP (version 2.1.11) (Price *et al.* 2010), then FigTree (version 1.4.4, https://github.com/rambaut/figtree/releases) was used to root the resulting tree on the edge between birds and *Bos taurus*. The alignment and rooted tree were then used as input by PAGAN (version 0.61) (Löytynoja *et al.* 2012) for the purpose of ancestral reconstruction. The “binary-table-by-edges-v2.1py” script was used to generate a table that reported upon whether a given position was mutated along each tree edge, and the “add-convention-to-binarytable-v.1.1.py” script was used to apply *Bos taurus* positional information to the output. The predicted ancestral and descendant values at the edge leading to hummingbirds were generated using the script “report-on-F-values-v1.1.py”, and all possible characters that could be found at each amino acid position of interest across the entire tree (including *Bos taurus*) were extracted by the script “extract-position-values_species_and_nodes-v1.1.py”. Scripts developed and used during this study can be found at https://github.com/corydunnlab/hummingbird.

### Modeling and simulation

We constructed small and large model systems of bovine mitochondrial cytochrome *c* oxidase from the high-resolution (1.5 Å) crystal structure (PDB 5B1A) (Yano *et al.* 2016). The large model system comprised all thirteen subunits, whereas in the small model only two catalytic subunits (COI and COII) were included, thereby allowing longer timescale simulations. Both wild-type and mutant (A153S) cases were considered within bovine model systems. The larger protein system was embedded in a multicomponent lipid bilayer (POPC:POPE:cardiolipin in 5:3:2 ratio) and only single component bilayer (POPC) was used in the case of two subunit complex, both constructed using CHARMM-GUI (Lee *et al.* 2016). Solvent water and Na^+^ and Cl^−^ ions (150 mM each) were added. In both setups, metal centers were in oxidized states with a water and an hydroxyl ligand at heme *a*_3_ and Cu_B_, respectively. Crosslinked Y244 was anionic [see also (Malkamäki and Sharma 2019)]. All amino acids were in their standard protonation states, except E242, K319 and D364 of COI, which were kept in a neutral state. The CHARMM force field parameters for protein, water, lipids and metal centers were taken from (MacKerell *et al.* 1998; Klauda *et al.* 2010; Best *et al.* 2012).

Additional subunit COI/COII homology models of hummingbird cytochrome *c* oxidase [both wild-type (S153) and mutant (A153) model systems] were constructed using bovine structure and the predicted amino acid sequence of *Florisuga mellivora* (accession YP_009155396.1). The MODELER program (Sali and Blundell 1993) was employed, using the default settings, to construct the homology models.

In order to test the reliability of our homology modeling procedures, we first modeled bovine subunit COI/COII sequences using the constructed homology model of hummingbird. Upon comparison of the output with the high resolution bovine COX crystal structure, we found RMS deviation of backbone atoms and sidechains to be ~0.3 Å and 1.57 Å, respectively. To consolidate our modeling protocol further, we performed similar analyses by comparing the human structure [PDB 5Z62, (Zong *et al.* 2018)] to the constructed bovine model and the bovine structure to the human homology model. For the catalytic subunit (91% sequence identity), this resulted in 0.67/0.68 Å and 1.68/1.88 Å RMSD for the backbone and sidechains, respectively. Overall, the lower values of RMSD values for comparison between models and structures suggest that our homology modeling procedure is quite robust.

All MD simulations were performed with GROMACS software (Abraham *et al.* 2015). Prior to production runs, all model systems were subjected to energy minimization followed by an equilibration MD. During equilibration, Nose-Hoover thermostat (Nosé 1984; Hoover 1985) and Berendsen barostat (Berendsen *et al.* 1984) were applied. LINCS (Hess 2008) algorithm implemented in GROMACS was applied to achieve the time step of 2 fs. During production runs, Nose-Hoover thermostat and Parinello-Rahman barostat (Parrinello and Rahman 1981) were applied to keep temperature and pressure constant at 310 K and 1 atm, respectively. The large and small model systems of bovine oxidase were simulated for 1.5 and 3 μs, respectively. The hummingbird COX wild-type and mutant models were simulated for 1 μs each, resulting in a total of 11 μs of atomistic simulations. VMD (Humphrey *et al.* 1996) was used for the visualization of trajectories and analysis.

## DATA AVAILABILITY

The authors affirm that all data necessary for confirming the conclusions of the article are present within the article, figures, and tables. Sequence information used for phylogenetic analysis can be found within the Dryad data repository (https://doi.org/10.5061/dryad.x69p8czf7).

## RESULTS AND DISCUSSION

### *Hummingbird mitochondrial DNA encodes an unusual cytochrome* c *oxidase subunit I substitution*

We sought specific coding changes within mtDNA-encoded genes that might be associated with the extreme metabolic capabilities of hummingbirds. Toward this goal, we used software with the capacity to predict ancestral sequences (Löytynoja *et al.* 2012) and software developed within the context of this study to identify all mtDNA-encoded amino acid positions mutated along the lineage leading to hummingbirds. Consistent with a link between mtDNA alterations and the unusual metabolic properties of these animals, the lineage leading to the family Trochilidae exhibits the greatest number of amino acid substitutions when considering 635 internal edges of a bird phylogenetic tree (Figure 1A). Of those 208 positions altered along the edge leading to hummingbirds (Table S1), the most conserved amino acid position mutated during establishment of hummingbirds was COI position 153 (Figure 1B; for convenience, we use the amino acid numbering associated with the structurally characterized *Bos taurus* COI subunit). This non-conservative A153S substitution was universal among all 15 hummingbird COI sequences obtained from the RefSeq (O’Leary *et al.* 2016) database (Table S2), yet was absent from all other birds upon examination of an alignment of 645 Aves COI entries (Figure 1C).

**Figure 1:**
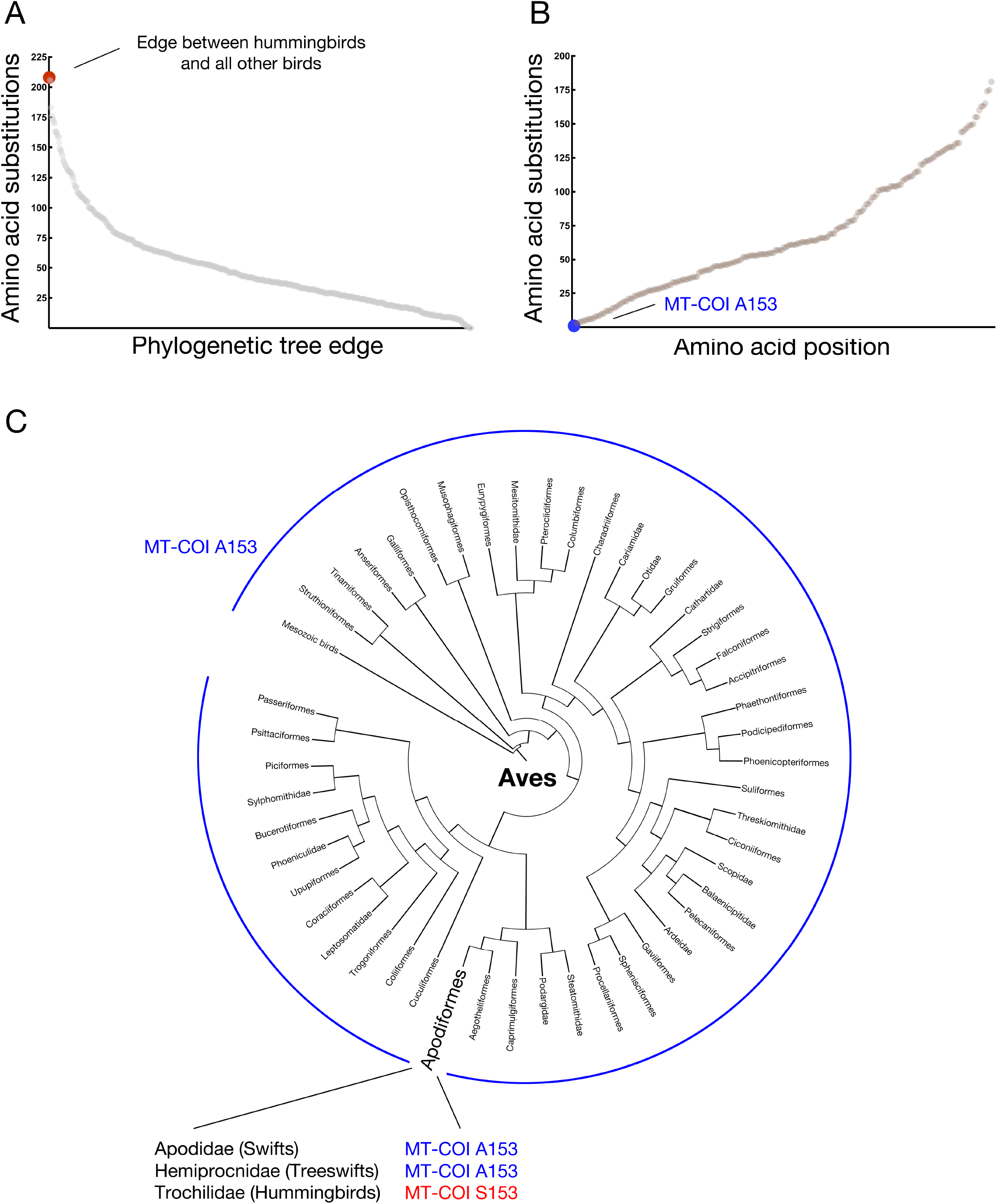
A rare alanine to serine substitution at bovine COI position 153 is universal among hummingbirds. (A) The edge leading to hummingbirds exhibits the largest number of substitutions within mitochondria-encoded proteins among all internal edges in a bird phylogenetic tree. A maximum likelihood tree was generated from an alignment of concatenated mitochondrial proteins from birds and *Bos taurus* using T-coffee in regressive mode (Garriga *et al.* 2019), followed by ancestral prediction using PAGAN (Löytynoja *et al.* 2012). Amino acid substitutions between each pair of ancestral and descendant nodes internal to the bird tree (node-to-node) were determined, summed across all positions, and plotted. (B) Among those changes found within the edge leading to hummingbirds, substitution at COI position 153 is most infrequent among birds, occurring only once. A plot demonstrating the number of times a given amino acid position was altered within the bird phylogeny is provided. (C) Serine at COI position 153 is unique to, and universal among, hummingbirds, as confirmed by phylogenetic analysis and by examination of an alignment of 645 Aves COI entries. Bird orders are arranged based upon a supertree modified from (Davis and Page 2014) under a Creative Commons license.

Since COI is the most commonly used DNA sequence barcode for studying animal ecology and speciation (Hill 2015; Pentinsaari *et al.* 2016), we next analyzed additional sequences covering the COI region of interest that we obtained from the Barcode of Life Data (BOLD) Systems (Ratnasingham and Hebert 2007). Initially, we focused upon sequences from the bird order Apodiformes, a taxon encompassing hummingbirds and swifts. 914 of 915 informative samples annotated as hummingbird-derived were found to carry a serine at position 153 of COI (Table S3). The remaining sample is mis-annotated as hummingbird, as determined by BLASTN analysis of its barcode (Altschul *et al.* 1990). In contrast, all 110 non-hummingbird Apodiformes samples harbored the ancestral A153. Extending our analysis to all informative bird barcodes, only 15/36,636 samples (< 0.1%) not annotated as hummingbird or its parental clade diverged from alanine at position 153. Assuming that these COI alterations were not the result of sequencing or annotation errors, we found that changes to A153 outside of hummingbirds were not fixed within each identified genus (Table S4). No other COI change appears universally encoded by hummingbird mtDNA, and position 153 does not contact a nucleus-encoded subunit, suggesting the lack of a single compensatory change that would lead to substitution neutrality (Blier *et al.* 2001). Codons for alanine and serine are separated by a distance of only one base pair alteration, suggesting that sequence-level constraints do not explain the singular fixation of the non-conservative A153S substitution in hummingbird COI. Since A153 is nearly universal among birds, yet appears to be substituted for serine in all hummingbirds, the A153S change within hummingbird COI may be linked to the aerobic capacity and prolonged hovering flight of these organisms.

### *Among vertebrates, the cytochrome* c *oxidase subunit I A153S substitution is fixed only in hummingbirds*

Beyond birds, substitution for alanine at COI position 153 was also extremely unusual among chordates, a taxon encompassing, and mostly represented by, vertebrates. Of 4,998 aligned Chordata sequences from the RefSeq dataset, only four non-hummingbird entries suggested a possible change at amino acid 153 (Table S5). Two RefSeq entries, from the sawtooth eel (*Serrivomer sector*) and the kuhli loach (*Pangio cf. anguillaris*), suggest the presence of the A153S substitution characteristic of hummingbirds. Another sequence, from the Lake Patzcuaro salamander (*Ambystoma dumerilii*), harbored a A153P substitution, and a COI entry from the air-breathing fish *Polypterus ornatipinnis* contained a A153V change. Subsequent analysis of accumulated COI barcodes from these genera suggested that any substitution at position 153 is not fixed or that the RefSeq entries are erroneous (Chen *et al.* 2017; Smith 2019). The apparent selective pressure against changes to COI position 153 among vertebrates, with the exception of the A153S substitution maintained within the family Trochilidae, is consistent with the hypothesis that this COX modification supports the extraordinary metabolism of hummingbirds.

### *Substitution of A153 within cytochrome* c *oxidase subunit I remains rare in metazoans, but has been fixed within several clades*

We proceeded to analyze metazoan COI sequences, and we found that substitution at A153 remains very rare. Indeed, only 146/7942 (< 2%) of informative RefSeq COI sequences report a substitution of A153 with any other amino acid (Table S6). In contrast to the results obtained upon analysis of vertebrate COI sequences, these changes appear common or fixed in a number of genera, as supported by analysis of BOLD sequence samples. Of note, a number of clams harbored changes at A153, including A153G, A153I, and A153S substitutions, which we speculate could be associated with the fluctuating oxygen concentrations commonly encountered by these molluscs (Ivanina *et al.* 2016). Of the clams encoding a A153G substitution, *Calyptogena magnifica* has previously been demonstrated to exhibit very low COX activity (Hand and Somero 1983), raising the possibility that G153 could be linked to diminished aerobic capacity within this clade. Moreover, a number of tube worms, including the hydrothermal vent resident *Riftia pachyptila*, harbored A153S substitutions. Tube worms, like some clams, are also subject to fluctuations in oxygen concentration and show resistance to periods of hypoxia (Arndt *et al.* 1998). However, *R. pachyptila* exhibits much higher COX activity than *C. magnifica* and maintains aerobic metabolism at low partial pressures of oxygen (Childress *et al.* 1984). Within the *Caenorhabditis* genus, some species, like *C. elegans,* encode an alanine at COI position 153, while others, including *C. afra*, encode serine at the same position (Table S6). Several nematodes, including *Caenorhabditis* worms, can synthesize either ubiquinone, associated with aerobic environments, or rhodoquinone, associated with anaerobic metabolism (Takamiya *et al.* 1999; Del Borrello *et al.* 2019; Tan *et al.* 2020). These findings suggest that these organisms also encounter fluctuating oxygen concentrations, prompting further speculation that changes to position 153 are linked to oxygen availability or consumption. The RefSeq sequences of a diverse set of additional organisms, including beetles, trematodes, myxozoans, and bees (discussed below), also indicated substitution from alanine at COI position 153.

Future efforts using worms found within the *Caenorhabditis* clade, which are eminently tractable for laboratory work, may provide experimental insight regarding the outcome of the A153S substitution. However, we note that it may be challenging to compare the effects of COI substitutions found and modeled within vertebrates to the effects potentially manifested in metazoans beyond vertebrates, as sequence divergence may limit potential interpretation of these identified substitutions. For example, while bovine and hummingbird COI, separated by ~310 million years (Kumar *et al.* 2017), are ~88% identical along the aligned region, bovine and bee COI, separated by more than 700 million years, are only ~66% identical, and bovine and nematode COI, also separated by greater than 700 million years, are only ~60% identical. Given that *Bos taurus* and the fungus *Saccharomyces cerevisiae* COI are 56% identical along the aligned region, we would be cautious about interpreting the changes we have identified here in the context of metazoan clades substantially divergent from vertebrates.

### Potential metabolic mimicry at position 153 of cytochrome c oxidase subunit I

As we considered the A153 substitutions that we identified among metazoans beyond the chordates, we noted the prominent presence of A153S, and the similar non-conservative substitution A153T, within several bee species. Analysis of BOLD samples from hymenopteran families Apidae and Megachilidae, the “long-tongued” bees (Cariveau *et al.* 2016), indicate nearly 100% substitution at COI position 153 to either serine or threonine (Table S7), while other families of bees harbor an ancestral alanine at position 153. Curiously, examination of COI sequences from millions of insect samples found in BOLD indicated that A153S and A153T conversion also characterizes many, but not all genera within the Eristalini tribe of dipteran hoverfly (Table S8). Intriguingly, adult hoverflies within this clade very closely mimic adult bees visually and behaviorally (Golding and Edmunds 2000) and, similar to adult bees, consume nectar and pollen (van Rijn *et al.* 2013). The identification of A153S and A153T substitutions in both bees and hoverflies hints at the exciting possibility of convergent evolution and metabolic mimicry, potentially rooted in diet and foraging behavior, at the mitochondria-encoded COI protein.

### Atomistic molecular dynamic simulations suggest that substitution at COI amino acid 153 may have functional consequences for proton transport in hummingbirds

To gain insight into the potential outcome of the A153S change encoded by hummingbird mitochondria, we focused our subsequent attention upon the crystal structure of a vertebrate COX. In the high-resolution (1.5 Å) crystal structure (Yano *et al.* 2016) of COX from *Bos taurus* (~ 88% identical in COI amino acid sequence to hummingbird along the aligned region), COI A153 is buried in the middle of COI transmembrane helix IV and is sandwiched between the D-channel, which is thought to conduct protons for oxygen reduction and PMF generation (Kaila *et al.* 2011; Wikström and Verkhovsky 2011), and a water-filled cavity lined with polar residues (Figure 2A). Moreover, residue 153 is only 10 Å from E242, a residue central to redox-coupled proton pumping by COX (Wikström *et al.* 2018; Wikström and Sharma 2018). These results, together with the high conservation at this position, suggested that alteration of A153 may affect enzyme activity.

**Figure 2:**
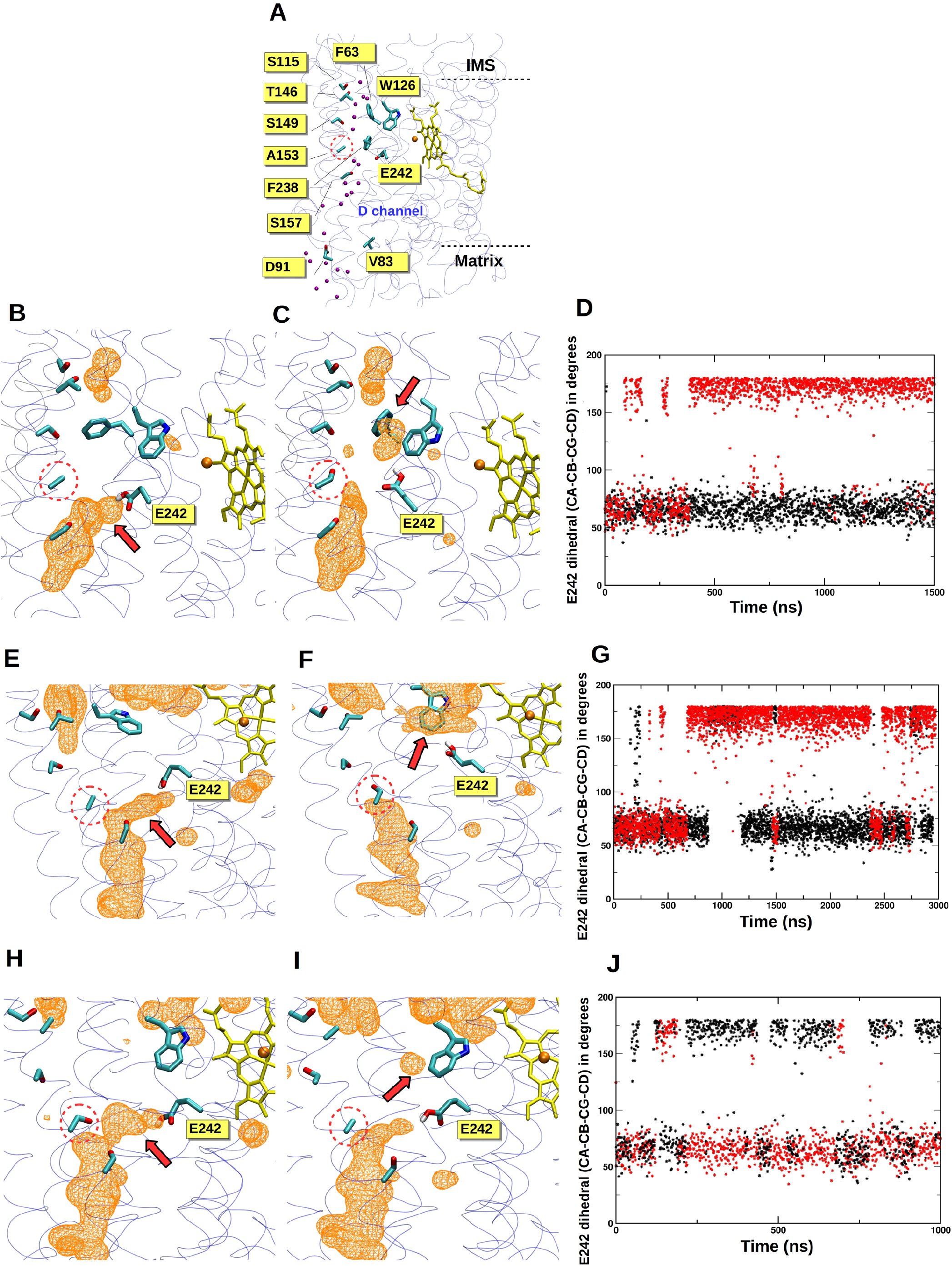
Hydration-coupled dynamics of conserved residue E242 are altered by the A153S substitution found in hummingbird COI. (A) The D-channel of proton transfer is located near residue 153 (dotted circles) in the high-resolution crystal structure of COX from *Bos taurus* (PDB 5B1A). Crystallographically resolved water molecules (purple spheres) in the domain above A153, together with nearby polar amino acids, form a potential proton transfer path. Cu_B_ is shown in orange and high spin heme in yellow. The catalytic COI subunit is shown with transparent ribbons and amino acids are displayed in atom-based coloring. (B-D) COI hydration and E242 side chain position are altered by substitution at COI position 153 in a large bovine COX simulation. (B) illustrates the native structure (A153) and (C) demonstrates the effects of A153S substitution. A red arrow highlights major changes to hydration, and water occupancy is shown as an orange colored mesh at an isovalue of 0.15. (D) E242 side chain dihedral angle (χ2) within COI encoding A153 (black) or S153 (red) during 1.5 μs of bovine large model simulation is displayed. Here, E242 adopts a predominant ‘up’ conformation within A153S substituted COI. (E-G), as in (B-D), but a small bovine model simulation has been deployed. (H-J), as in (B-D), but a small hummingbird model has been simulated. In (J), S153 (red) is wild-type and A153 (black) is mutant.

To further investigate the functional relevance of the A153S change in hummingbirds, we performed atomistic classical MD simulations on two different vertebrate model systems at differing scales. Remarkably, multiple microsecond simulations demonstrated changes in hydration within the vicinity of position 153 that were coupled to the dynamics of the aforementioned E242. Specifically, during simulations of the entire 13-subunit wild-type bovine COX performed in membrane-solvent environment, E242 was typically found in the ‘down’ state (χ2 ~ 60°), extending towards the D-channel proton uptake site D91 (Figures 2B and 2D). In contrast, upon A153S substitution, the bovine E242 commonly swung to the ‘up’ position (χ2 ~ 180°, Figures 2C and 2D). Similar findings emerged (Figures 2E-G) from longer simulations performed on small bovine model systems, suggesting that the observed behavior is robust. The microscopic changes in hydration near E242 stabilized its ‘up’ position (Figures 2C and 2F) and resulted in its connection to the COI regions near the positively charged intermembrane space via water molecules (Figure S1).

Next, we performed simulations using a constructed hummingbird homology model based upon the primary amino acid sequence of the hummingbird *Florisuga mellivora*. Our methodology for hummingbird homology model construction was validated by comparison of bovine and human COI homology models with existing structures, which yielded low root-mean-square deviation values (see MATERIALS AND METHODS). E242 behavior and channel hydration was again dependent upon whether alanine or serine was present at position 153 (Figure 2H-I), although the effect was less prominent than in bovine models. Here, in the constructed hummingbird model containing its wild-type S153 variant, E242 was stabilized in the ‘down’ position. Upon S153A replacement, both ‘up’ and ‘down’ populations were observed, and increased motility was visualized (Figure 2J) with corresponding changes to local hydration (Figure 2I). Together, our results suggest that position 153, substituted for a polar serine in hummingbirds, can determine the behavior of the key catalytic residue E242.

Interestingly, our simulations suggest that the behavior of additional amino acids beyond E242 may be affected by the amino acid found at position 153. For example, our hummingbird simulation strongly indicated that a change in F238 side chain angle is linked to E242 motion (Figure S2) and is influenced by whether residue 153 is an alanine or a serine. These data are supported by a GREMLIN co-evolution analysis (Kamisetty *et al.* 2013), initiated by use of the bovine COI sequence, that suggests co-evolutionary coupling between F238 and E242 (Table S9).

F238 has been suggested to play a key role in oxygen diffusion through COX (Mahinthichaichan *et al.* 2018), and indeed hummingbirds are characterized by a profound oxygen consumption rate during flight (Lasiewski 1962; Suarez *et al.* 1991; Clark and Dudley 2010; Fernández *et al.* 2011). Therefore, altered behavior of F238 upon A153S substitution prompted us to consider the possibility that oxygen access to COX is augmented in the hummingbird. However, we are equivocal regarding a scenario in which improved oxygen access is prompted by A153S substitution. First, with caveats related to the evolutionary divergence between bacteria and vertebrates, a S153A substitution in bacterial COI (bacterial A-family cytochrome *c* oxidases harbor a serine at this position within their catalytic subunit) led to similar cytochrome *c* oxidation rates and initial proton pumping rates (Pfitzner *et al.* 2000; Namslauer *et al.* 2007). Moreover, although *in vitro* assays may not reflect respiration rates obtained *in vivo* (Schwerzmann *et al.* 1989; Suarez *et al.* 1991), the oxygen consumption rate of isolated hummingbird mitochondria, when normalized to mitochondrial inner membrane area, did not notably differ from mammalian mitochondria (Suarez *et al.* 1991). Nevertheless, the tight evolutionary as well as dynamic coupling between F238 and E242 is likely of functional importance. For instance, the F238 flip (also E242) in hummingbird COI causes partial hydration in the region (Figure S2B), which would impede oxygen availability to the active site. Such features may play an important role in fine tuning overall respiration rates in a manner dependent upon external factors.

### An additional COI variant ancestral for hummingbirds and rare among other birds

One other COI variant beyond S153 appeared restricted to hummingbirds (Tables S1) when considering bird sequences obtained from the RefSeq database. Among the 15 hummingbird COI sequences found within this dataset, nine contained a conservative V83I substitution that is found in no other bird entry (Table S2). Expanding our analysis to Apodiformes barcodes obtained from BOLD, 110/110 non-hummingbird samples carried the V83 allele ancestral for birds. In contrast, 671/929 informative hummingbird samples within this dataset carried a V83I substitution, and 258/929 samples harbored a valine at this position (Table S3). Looking more broadly at 36,878 bird samples, substitution at V83 is extremely rare among birds (< 0.1%), although unlike the A153S substitution, this V83I allele may be widely shared among members of a limited number of non-hummingbird genera (Table S4). Phylogenetic analyses based upon a tree of mtDNAs obtained from RefSeq (Figure S3) or from the BOLD database (not shown) suggest that the V83I substitution was present at the establishment of hummingbirds, then subsequently reverted to valine several times during expansion of this clade. Within the COX enzyme, amino acid 83 lies within 9 Å of COI residue D91, which contributes to proton uptake via the eponymous D-channel described above (Figure 2A). Position 83 is also located within 6 Å of N80, a component of the ‘asparagine gate’ which is thought to move protons toward the active site (Henry *et al.* 2009; Liang *et al.* 2017; Ghane *et al.* 2018).

### What is the phenotypic outcome of these hummingbird COI substitutions?

Raising the possibility of altered proton movement at hummingbird COX, our results show clear changes to the behavior of key D-channel residues and to surrounding hydration when comparing models swapping alanine for serine, or vice versa, at COI position 153. Interestingly, previous studies of bacterial respiration suggest that amino acid 153 can influence coupling between electron transport and COX proton pumping, further indicating that proton motility may have been a focus of selection during evolution of hummingbirds. Specifically, a serine to aspartic acid change made at the analogous amino acid position in *Rhodobacter sphaeroides* COI abolished proton pumping while allowing electron transport coupled to proton transfer from the periplasmic side of the bacterial inner membrane when the enzyme was analyzed under zero PMF conditions (Pawate *et al.* 2002; Namslauer *et al.* 2007). Further suggesting unusual proton handling by hummingbird COX, the V83I substitution ancestral to hummingbirds is located near the ‘asparagine gate’ at the matrix side of the mitochondrial inner membrane, and mutations near this site lead to changes in the number of protons pumped per oxygen reduction (Pfitzner *et al.* 2000). Also of note, functional links have been suggested to exist (Vakkasoglu *et al.* 2006) between the asparagine gate and the key E242 residue, the behavior of which is clearly affected by A153S mutation.

We suggest two potential outcomes of the D-channel changes prompted by the A153S change universal to hummingbirds. First, if the bovine models accurately reflect the outcome of this substitution, hydration differences associated with the presence of a polar residue at position 153 may promote intrinsic uncoupling (Murphy and Brand 1988) of COX when the PMF is high across the mitochondrial inner membrane, even leading to the use of protons from the intermembrane space for oxygen reduction under conditions of high polarization (Mills *et al.* 2002). Rapid, on-site decoupling of proton pumping from electron transport may serve as a local response to cessation of flight, when an immediate rise in matrix ATP might lead to excessive PMF, ETC over-reduction, and reactive oxygen species (ROS) production (Papa *et al.* 1997; Kadenbach 2003). Intrinsic, local, and immediate uncoupling might be particularly important within hummingbird muscle, where the densely packed ETC components might generate a destructive wave of ROS linked to inner membrane hyperpolarization. Proton conductance across the hummingbird mitochondrial inner membrane has not, to our knowledge, been assessed under multiple mitochondrial membrane potentials. Such an approach could reveal enzyme uncoupling linked to membrane hyperpolarization (Kadenbach 2003).

Second, hummingbirds require robust thermoregulation, as these organisms exhibit high surface to mass ratios, are found at elevations associated with low temperatures, and are subject to convective heat loss while engaged in hovering flight (Altshuler and Dudley 2002; Altshuler *et al.* 2004; Projecto-Garcia *et al.* 2013). Beyond potential heat generation promoted by COX ‘slip’, or decoupling of electron transport from proton pumping (Musser and Chan 1995), results emerging from our bovine models raise the possibility that changes to COI hydration accompanying A153S substitution might allow direct, albeit limited movement of protons across the inner membrane that could contribute to non-shivering thermogenesis. Interestingly, thermoregulation may act as an initial selective force toward increased metabolic capacity (Bennett 1991) and therefore may have played a particularly prominent role during the evolution of hummingbirds.

Thus far, the vertebrate mitochondrial genome remains refractory to directed modification toward a desired sequence change (Patananan *et al.* 2016), preventing a direct test of the hummingbird-enriched COI substitutions in the background of a hummingbird mtDNA or of a related, non-hummingbird mitochondrial genome. However, we hope that future biochemical experiments guided by our combined use of phylogenetic analysis and atomistic simulations, may be informative regarding the role of hummingbird COI changes in these organisms. Excitingly, other changes to mtDNA-encoded oxidative phosphorylation machinery beyond COX are rare among other birds yet common in hummingbirds, and these substitutions await further analysis. Finally, while mtDNA sequence is far more prevalent, we expect that accumulating nuclear DNA sequence information from birds (Zhang *et al.* 2015) will allow analysis of divergent, nucleus-encoded members of the oxidative phosphorylation machinery and their potential role in hummingbird metabolism.

## ACKNOWLEDGEMENTS

C.D.D. is funded by an ERC Starter Grant (RevMito 637649) and by the Sigrid Jusélius Foundation. V.S. is funded by the Academy of Finland, the University of Helsinki, and the Sigrid Jusélius Foundation. We thank Fyodor Kondrashov for advice on COI sequence recovery and alignment, Tamara Somborac for comments on the manuscript, Gregor Habeck for assistance in use of R Studio, Aapo Malkamäki for support in MD simulation setup, and Mårten Wikström for helpful discussions. We also acknowledge the Center for Scientific Computing, Finland for their generous computational support.

## AUTHOR CONTRIBUTIONS

C.D.D. and B.A.A. performed phylogenetic and taxonomic analyses. V.S. performed molecular dynamics simulations. All authors prepared the manuscript text and figures.

## SUPPLEMENTARY INFORMATION

**Figure S1: Water-based paths toward the positively charged intermembrane space.** Due to shift in ‘down’ to ‘up’ conformation of E242, the side chain of this amino acid connects to the hydrophilic region above heme propionates and near amino acids Y231 and T146 following A153S substitution in a small bovine COI simulation.

**Figure S2: The position of the COI F238 side chain is coupled to E242 dynamics.** (A) F238 is shown within the hummingbird small model simulation when E242 is in the (A) down position and when E242 points in the (B) up position. F238 dihedral angle (χ1, N-CA-CB-CG) is compared to E242 dihedral angle (χ2, CA-CB-CG-CD) in the (C) bovine large model simulation and in the (D) hummingbird small model simulation. E242 data from Figures 2D and 2J are shown, with F238 dihedral angle for the A153 variant plotted in blue and for the S153 variant plotted in orange.

**Figure S3: COI I83 is ancestral for hummingbirds and reverts to the bird consensus V83 along multiple lineages.** Mitochondrial genome sequences (RefSeq release 91) of the indicated organisms were obtained (NC_008540.1, NC_034933.1, NC_010094.1, NC_030286.1, NC_024156.1, NC_033406.1, NC_033404.1, NC_033405.1, NC_027453.1, NC_027454.1, NC_030285.1, NC_033414.1, NC_025786.1, NC_027455.1, NC_030287.1, NC_030288.1, NC_033413.1) and aligned by MAFFT using the G-INS-i iterative refinement method. Next, a phylogenetic analysis was performed within the MEGAX software (Kumar *et al.* 2018; Stecher *et al.* 2020) using a maximum likelihood approach based upon the Tamura-Nei model (Tamura and Nei 1993). A gamma distribution of six categories was used for modeling substitution rate differences between sites, with some sites potentially invariant. Positions with less than 90% coverage were removed. We display the tree with the greatest log likelihood value (−98938.63) with the results of 100 bootstrap replications noted at each node. The amino acid found at COI position 83 is listed next to each organism within the resulting tree.

**Table S1: Analysis of the frequency at which mutations occur at specific positions mutated along the lineage to hummingbirds.** Following concatenation and alignment of protein sequences encoded by birds and by *Bos taurus*, a maximum-likelihood tree was generated, and 208 positions harboring substitutions that occurred along the lineage to hummingbirds were predicted. The total number of times each position was found to mutate within birds is provided, along with the predicted ancestral and descendant amino acids on the edge leading to hummingbirds, and all amino acid possibilities at that position (including within *Bos taurus*). Whether substitutions identified within the RefSeq database at a given position are limited only to edges either within or leading to the hummingbird clade is also noted.

**Table S2: Analysis of full-length bird COI sequences obtained from the RefSeq database.** Sequences of all mitochondria-encoded proteins found in the NCBI RefSeq database (release 92) (O’Leary *et al.* 2016) were downloaded, and the COI FASTA sequences of 645 birds were extracted. MAFFT alignment was performed (Katoh and Standley 2013) using the G-INS-i iterative refinement approach, and those birds with a A153S substitution are listed along with the variant harbored by each species at COI position 83.

**Table S3: Examination of amino acids 83 and 153 in Apodiformes barcodes obtained from BOLD.** The query “Apodiformes” was made using the BOLD server (Ratnasingham and Hebert 2007) to recover COI FASTA sequences. MAFFT alignment was carried out using L-INS-i iterative refinement method and translated in AliView (Larsson 2014) using the vertebrate mtDNA translation table. Informative samples with annotated species names were assessed to determine the amino acids found at positions 83 and 153.

**Table S4: Analysis of Aves COI barcodes obtained from BOLD.** The query “Aves” was made using the BOLD server (Ratnasingham and Hebert 2007) to recover COI FASTA sequences. MAFFT alignment was carried out under the “auto” setting and translated in AliView (Larsson 2014) using the vertebrate mtDNA translation table. Informative COI samples outside of the order Apodiformes and not harboring alanine at position 153 or valine at position 83 are listed.

**Table S5: Analysis of chordate COI sequences obtained from the RefSeq database.** COI sequences obtained from the RefSeq database (O’Leary *et al.* 2016) were filtered for chordate entries. MAFFT alignment was performed using the “auto” setting and poorly aligned sequences were deleted. Hummingbird sequences were removed. Four informative, non-hummingbird sequences were characterized by substitution of A153 for other amino acids, although analysis of annotated BOLD accessions labelled with a species name or entry suggest that these COI substitutions are not generally shared throughout each genus.

**Table S6: Analysis of metazoan COI sequences obtained from the RefSeq database.** COI sequences obtained from the RefSeq database (O’Leary *et al.* 2016) were filtered for metazoan entries. Analysis was performed as in Table S5, and non-chordate samples with a substitution at A153 are displayed. For those species for which RefSeq suggests a substitution at this position, BOLD entries for each genus of the COI-5P class were aligned using MAFFT under the FFT-NS-2 option and translated in AliView (Larsson 2014) using the indicated mtDNA translation table. Informative entries harboring A153 or substitutions from the ancestral amino acid are quantified.

**Table S7: Analysis of bee COI barcodes obtained from BOLD.** Entries for bee families Apidae, Megachilidae, Colletidae, Halictidae, Andrenidae, and Melittidae, as defined in (Sann *et al.* 2018), were obtained from BOLD (Ratnasingham and Hebert 2007), but no samples from the small bee family Stenotritidae were available. In addition, no Spheciformes samples were found in BOLD. After deletion of samples not of the COI-5P class or not associated with a species, MAFFT alignment was carried out under the “auto” setting. DNA sequences were translated to the informative reading frame in AliView (Larsson 2014) using the invertebrate mtDNA translation table. Substitution frequencies among each family is provided, along with the variant found at position 153 in each sample.

**Table S8: Analysis of COI barcodes from Eristalini tribe of hoverflies obtained from BOLD.** Entries for the Anasimyia, Eristalinus, Eristalis, Helophilus, Lejops, Mesembrius, Palpada, Parhelophilus, Phytomia, Senaspis, Mallota, and Chasmomma genera were obtained and analyzed as in Table S7.

**Table S9: GREMLIN analysis of co-evolution.** GREMLIN (Kamisetty *et al.* 2013) was run using the default settings and the bovine COI sequence as a query. ‘i_id’ and ‘j_id’ represent analyzed residues, ‘r-sco’ represents the raw scoring, ‘s-sco’ represents the scaled score generated from the raw score and average of the raw score, and ‘prob’ represents the probability of residue contact given the scaled score and the sequences per length.

