## Supplementary figures and images for "An unusual amino acid substitution within hummingbird cytochrome *c* oxidase alters a key proton-conducting channel"

### Dunn_et_al_Hummingbird_Figure_S1.tif

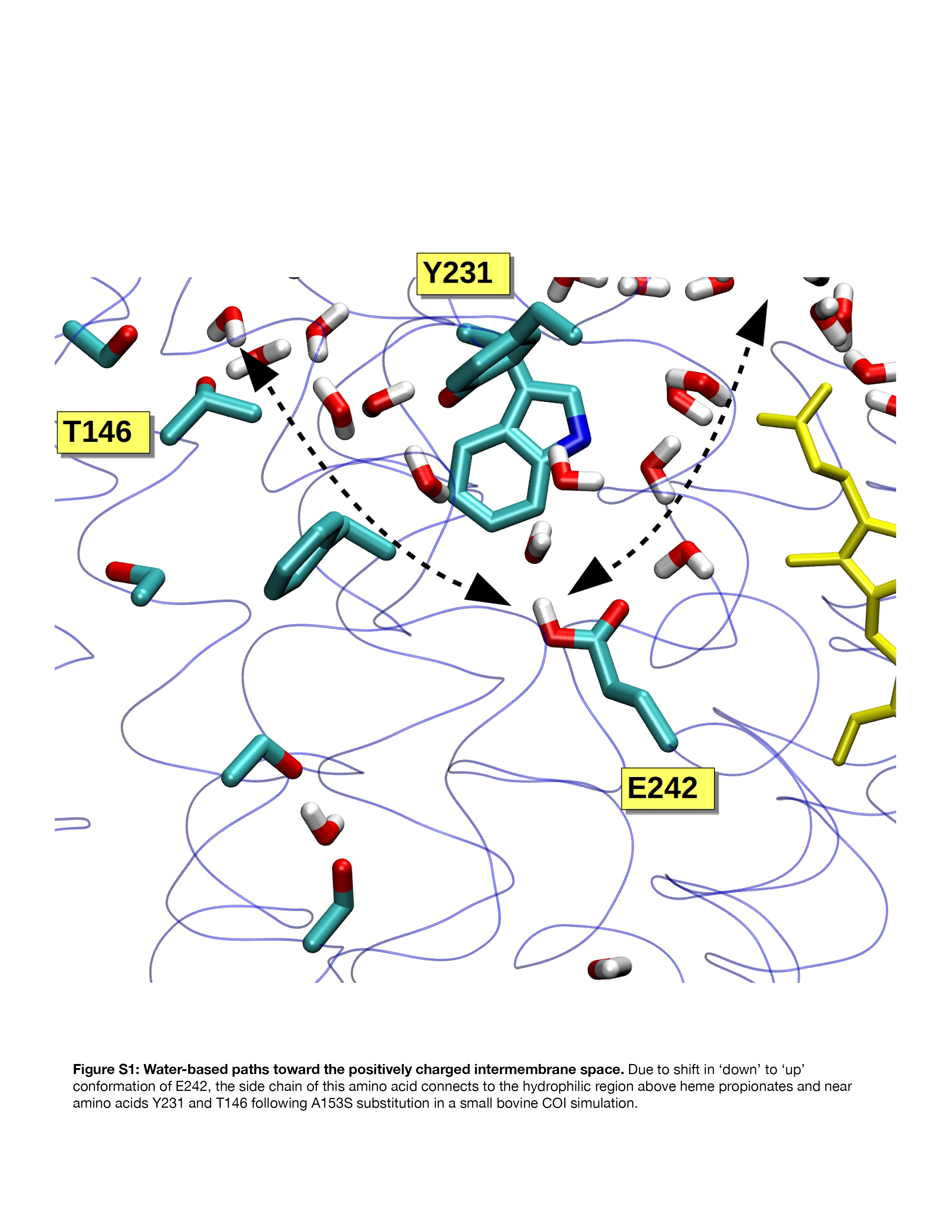

### Dunn_et_al_Hummingbird_Figure_S2.tif

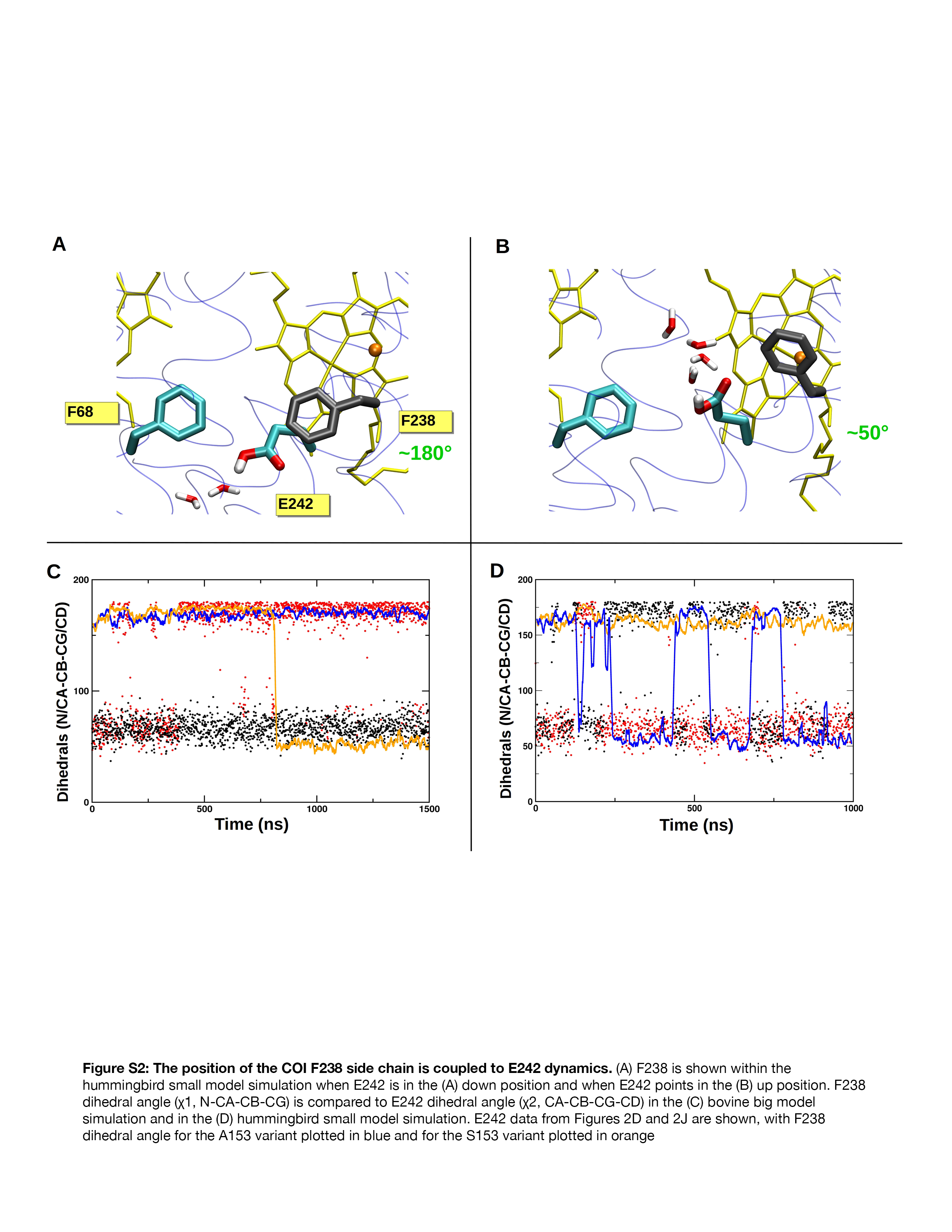

### Dunn_et_al_Hummingbird_Figure_S3.tif

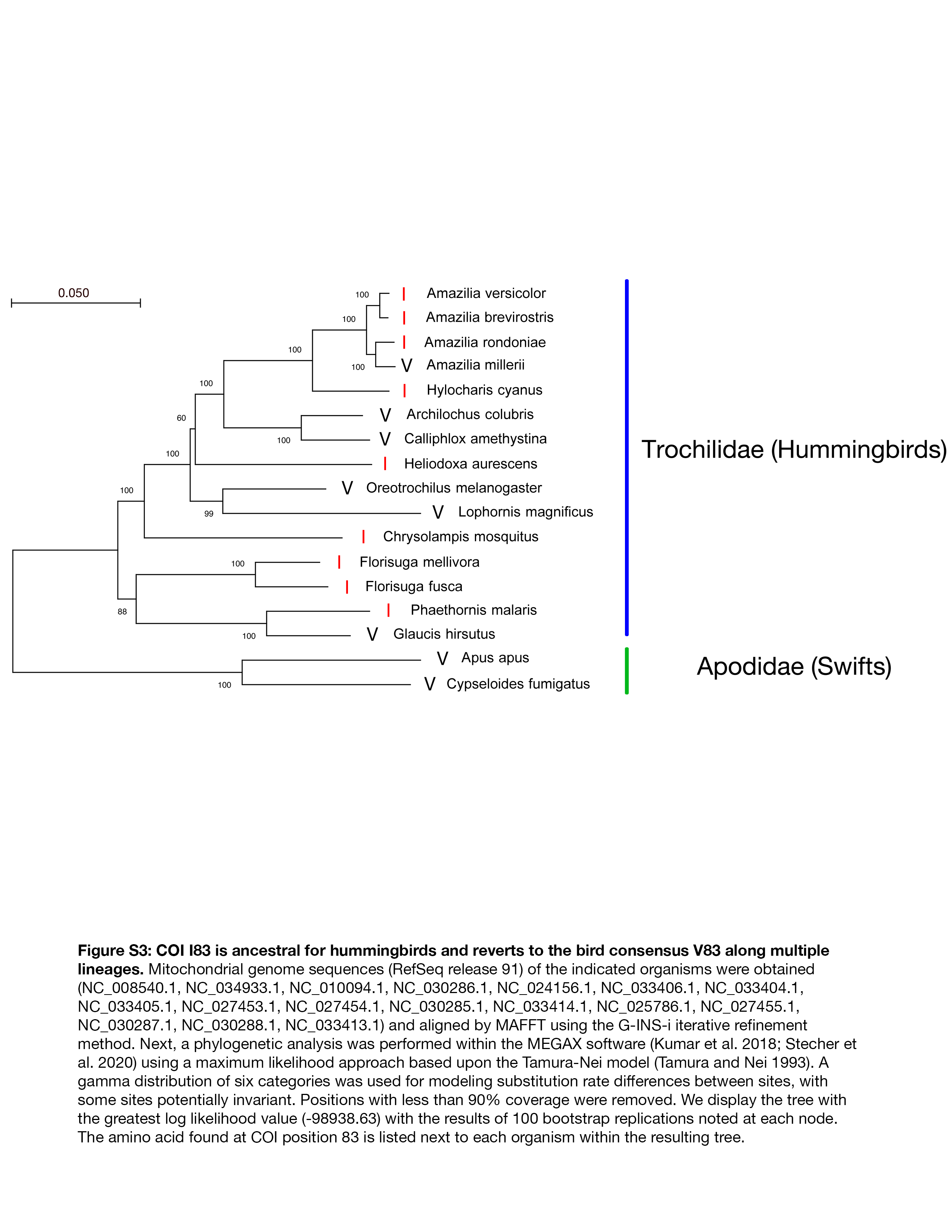
